# Glutathione overproduction mediates lymphoma initiating cells survival and has a sex-dependent effect on lymphomagenesis

**DOI:** 10.1101/2023.11.22.568023

**Authors:** H.-Alcántara Alberto, Omar Kourani, Ana Marcos-Jiménez, Patricia Martínez-Núñez, Estela Herranz-Martín, Patricia Fuentes, María Luisa Toribio, Cecilia Muñoz-Calleja, Teresa Iglesias, Miguel R. Campanero

**Author notes:** **Corresponding author:** Miguel R. Campanero. AH-A and OK contributed equally to this work.

## Abstract

Lymphoid tumor patients often exhibit resistance to standard therapies or experience rapid relapse post-remission. Tumor-initiating cells (TICs), a small fraction of the tumor cell population known for their self-renewal capacity and resistance to cancer therapies, likely drive tumor relapse. Tumorigenicity strongly correlates with growth in soft gels and TICs are the only cancer cells capable of growing in soft gels. Targeting pathways critical for TIC survival or growth holds promise for improving cancer treatment outcomes but TIC biology remains poorly understood. Here, we show that culturing lymphoid cells in soft hydrogels triggers reactive oxygen species (ROS) production, leading to non-tumor lymphoid cell death while enabling the survival and proliferation of a subset of lymphoma/leukemia cells, TICs or TIC-like cells. Treatment with the antioxidant N-acetylcysteine inhibits this lethality and even promotes the growth of primary non-tumor lymphoid cells in soft gels. Some lymphoma cells escape ROS-induced lethality by boosting antioxidant glutathione production, a response not seen in non-tumor cells. Reducing glutathione production in lymphoma cells, either through pharmacological inhibition of glutamate cysteine ligase (GCL), the enzyme catalyzing the rate-limiting step in glutathione biosynthesis, or via knockdown of *GCLC*, the GCL catalytic subunit, sharply decreased cell viability and proliferation in soft gels and tumor growth in immunodeficient mice. Tumor cells from B-cell lymphoma/leukemia patients and λ-MYC mice, a B-cell lymphoma mouse model, overproduce glutathione. Importantly, pharmacological GCL inhibition hindered lymphoma growth in female λ-MYC mice, suggesting that this treatment holds promise as a therapeutic strategy for female lymphoma/leukemia patients.

## Introduction

Many conventional cancer therapies often lead to undesired side effects because they target proteins and cellular processes necessary for normal cell survival and proliferation, including various progenitor cell types. Identifying genes and proteins specifically associated with tumor traits would facilitate the development of therapies that selectively target tumor cells while minimizing toxic effects. Despite some potentially fatal side effects, the major failures of current cancer treatments are the lack of response in certain human cancer types and tumor relapse after initial remission^1^.

Despite progress in cancer research and the development of new therapies, many cancer types still have limited treatment options. Indeed, one-third of cancer patients succumb to the disease within five years of diagnosis, primarily due to tumor recurrence^1,2^. Discovering strategies to prevent tumor relapse continues to be a challenge.

Growing evidence suggests that numerous cancers, including hematopoietic tumors, may be driven by a small subset of cells known as tumor-initiating cells (TICs)^3^. These TICs exhibit unique properties such as self-renewal and high tumorigenicity, making them the only cancer cells capable of regenerating the original tumor^4^. Since these cells are the most resistant cells to chemotherapy and radiotherapy within the tumor^5^, they are considered as responsible for tumor relapse^3^. Their resistance has been linked to their capacity to accumulate intracellular antioxidants, reducing reactive oxygen species (ROS) levels^6,7^.

While the discovery of therapeutic strategies targeting TICs holds promise, our understanding of TIC biology remains limited due to the challenges in identifying and isolating them. Conventional cell surface markers are often unreliable and vary among cancer types^8,9^. However, a breakthrough came with the observation that TICs, rather than the bulk of differentiated tumor cells, thrive in soft hydrogels^10,11^, offering an opportunity to uncover signaling pathways specific to TIC growth and identify TIC-specific therapeutic targets.

Growth in soft agar hydrogels strongly correlates with tumorigenicity^12^, and this property has long been used to distinguish tumorigenic from non-tumorigenic cells *in vitro*. Carcinoma cells, for instance, form colonies in soft agar hydrogels, whereas immortalized NIH-3T3 fibroblasts do not^13–15^. Similarly, lymphoblastoid B-cell lines (LCLs), non-tumor cell lines created through the immortalization of normal B-cells from healthy donors using the Epstein-Barr virus (EBV), do not grow in soft agar gels. In contrast, a small fraction of lymphoma and leukemia cell lines successfully grow in these gels^16,17^, suggesting the presence of TIC or TIC-like cells even in established lymphoma/leukemia cell lines cultured *in vitro*. These findings align with previous reports indicating the maintenance of stem-cell-like cancer cells in established carcinoma cell lines^18^.

It has been proposed that epithelial and mesenchymal cells survival relies on the signaling provided by integrins attached to a rigid surface and that culturing them in soft gels leads to cell death by preventing attachment to a rigid surface^13^. Importantly, soft-agar hydrogels not only prevent anchorage to a rigid surface but also to other cells. It should be noted that while lymphoid cells remain viable in the circulatory system without anchoring to a rigid surface, they do not proliferate there. Instead, they proliferate in lymphoid organs, where interactions with extracellular matrix, stromal cells, and/or other lymphocytes may play a role. This suggests that the proliferation of non-tumor lymphoid cells may also depend on anchorage.

Despite the strong correlation between growth in soft gels and tumorigenicity^12^, it remains unclear why TICs are the exclusive tumor cells capable of growing in soft gels, while non-tumor lymphoid cells do not. Investigating the molecular mechanisms enabling TIC growth in soft gels could unveil new targets to selectively inhibit TIC growth and prevent lymphomagenesis. To address this, we have established experimental conditions for retrieving cells from soft agar and analyzing them by flow cytometry. These studies indicate that overproduction of glutathione drives the survival and growth of a small subset of lymphoid tumor cells in soft gels and xenografts, likely TIC or TIC-like cells, and that glutathione is selectively required for tumor growth in females of a mouse model of lymphoma.

## Methods

### Cell lines and human samples

Human Burkitt lymphoma DG-75 (CRL-2625) and BL-2 (ACC-625), Diffuse Large B-Cell lymphoma Toledo (CRL-2631), T-cell leukemia Jurkat (Clone E6-1, TIB-152), and HEK-293T (CRL-1573) cell lines were obtained from ATCC (LGC Standards S.L.U., Barcelona, Spain). LCLs JY and X50-7 were from the European Collection of Authenticated Cell Cultures (ECACC 94022533) and Cellosaurus (RRID:CVCL_8277), respectively. DG-75, BL-2, Toledo, and Jurkat were cultured in RPMI 1640 medium (21875034), whereas HEK-293T were cultured in Dulbecco’s modified Eagle’s medium (DMEM) (11965092), both from ThermoFisher Scientific (Waltham, MA, USA). All cells were mycoplasma-negative. Both media were supplemented with 10% heat-inactivated fetal bovine serum (FBS) and 2 mM glutamine (25030081), all from ThermoFisher Scientific, plus 100 units/mL penicillin (Laboratorios ERN.S.A., Madrid, Spain; 804443) and 100 μg/mL streptomycin (Reig Jofre S.A., Madrid, Spain; 753483) and were maintained at 37°C in a humidified incubator with 5% CO_2_.

Human peripheral blood lymphocytes (PBLs) of healthy donors (HD) were isolated as described ^19^ from buffy coats, obtained from the Madrid Blood Donor Centre (Madrid, Spain), and then stimulated with 5 µg/ml leucoagglutinin (L4144; Sigma-Aldrich, Madrid, Spain) and cultured in RPMI 1640 medium supplemented with 10% heat-inactivated FBS, penicillin, streptomycin, L-glutamine, and 50 U/mL of human recombinant interleukin 2 (IL2) ^20^.

Patients included in this study were diagnosed according to WHO and refined consensus criteria ^21,22^. Informed consent was obtained in accordance with the Declaration of Helsinki. Experimental procedures were approved by the Institutional Board of Hospital de La Princesa (PI-802). Cells isolation from freshly donated peripheral blood was done using Ficoll-paque plus density gradient centrifugation (Amersham Biosciences, Little Chalfont, UK). Peripheral blood mononuclear cells (PBMCs) from healthy donors, obtained from peripheral blood or buffy coats, were used as control. Cells were cultured in RPMI-1640 medium supplemented with 10% heat-inactivated FBS, L-glutamine, penicillin, and streptomycin at 37 °C in 5% CO2.

### Lentivirus production and cell transduction

Lentiviral particles were produced as previously described ^23^ employing, psPAX2 and pMD2G-VSVG plasmids [provided by D. Trono (Ecole Polytechnique Federale de Lausanne, Lausanne, Swisse)] and MISSION pLKO.1-puro-based vectors (Sigma-Aldrich, Madrid, Spain,) encoding either a non-targeting shRNA (SHC002) or *GCLC*-targeting shRNAs sh-62 (TRCN0000344862), sh-65 (TRCN0000333565), and sh-86 (TRCN0000048486). Conditioned medium was harvested, filtered through 0.45-μm filters, concentrated by ultracentrifugation (81550 g, 2h at 4°C) and stored at -80°C. DG-75 and Toledo cell lines were incubated with viral particles in culture medium with 8μg/ml protamine sulfate during 16h. Cells were washed to remove viral particles and transduced cells were selected in the presence of 1μg/ml puromycin (Sigma-Aldrich, P8833) for at least 96 h.

### Western Blotting

Cells were harvested and suspended in Total Lysis Buffer (0.125 M Tris-HCl pH 6.8, 4% SDS and 20% glycerol). Cell lysates were boiled for 15 min and protein concentration was determined by the Lowry method. After quantification, β-mercaptoethanol (1:100 v/v) and bromophenol blue powder were added. Protein samples were resolved in 8% SDS-PAGE and transferred to nitrocellulose membranes as described (6). Membranes were probed with anti-GCLC sc-390811 (200 ng/ml) and anti-Vinculin sc-73614 (200 ng/ml), both from Santa Cruz Biotechonology (Santa Cruz, CA, USA) followed by goat anti-mouse IgG conjugated to IR-800Dye (926-32210; 1:15,000) from LI-COR Biosciences (Lincoln, Nebraska, USA), and scanned using an Odyssey® Infrared imaging system (Model 9120, LI-COR Biosciences).

### Mouse lymphoma model and tumor xenografts

The previously described mouse model of Burkitt lymphoma C57BL/6N-Tg(IGL-MYC)3Hm (λ-MYC mice) ^25^ was obtained from the NCI mouse repository (Strain code 01XA7), backcrossed to C57BL/6J for more than 10 generations, and maintained in heterozygosity in a C57BL/6J background. Mice were monitored weekly for palpable tumors, commencing at an age of 8 weeks. Males and females were treated with 20 mM BSO (19176, Sigma-Aldrich) in the drinking water starting at 21 days of age, immediately after weaning. The experimental endpoint was 250 days of age unless palpable tumors were detected. At endpoint, mice were sacrificed and the spleen and all tumors from each individual mouse were resected and weighed.

Transduced DG-75 cells (2 x 10^6^) in 0.1 ml of phosphate buffer saline (PBS) were injected subcutaneously in the dorsal flanks of 8-to 10-week-old female NOD.CB17-Prkdcscid/J mice (Charles River Laboratories, Wilmington, MA, USA). All mice were inoculated with control cells in one flank and *GCLC*-silenced cells in the opposite flank. Tumor masses were removed after 3 weeks and weighted. All animal procedures were approved by the institutional review board. All animal procedures were approved by the CSIC Ethics Committee (ref. 634/2017 and 1053/2021) and by the Madrid Regional authorities (ref. PROEX 215/17 and 093.7/21), and conformed to EU Directive 2010/63EU and Recommendation 2007/526/EC regarding the protection of animals used for experimental and other scientific purposes, enforced in Spanish law under Real Decreto 1201/2005. Overall mouse health was assessed by daily inspection for signs of discomfort, weight loss, or changes in behaviour, mobility, and feeding or drinking habits.

### Transformation assays *in vitro*

Agar was prepared in RPMI medium supplemented with 10% heat-inactivated FBS, L-glutamine, penicillin and streptomycin. For colony formation assays, cells (5 x10^4^) were suspended in 2 mL of 0.33% noble agar (Difco) and laid over 6-well culture plates previously coated with 2 mL of 0.5% noble agar. When appropriate, soft agar was supplemented with 50 µM BSO. Plates were kept at 37°C in a humidified incubator in the presence of 5% CO_2_. The number of colonies formed after 3 weeks was counted in triplicate plates.

### Flow cytometry

Cell cycle analysis of cells cultured in liquid medium was performed as described ^26^ using a FACS Canto II flow cytometer (BD Bioscienes, San Jose, CA, USA). For cell cycle analysis of cells embedded in soft agar, cells (10^5^) were suspended in 1 mL of 0.33% noble agar, seeded in the bottom of 15-mL polypropylene tubes, cooled for 5 minutes on ice, and then kept at 37°C in a humidified incubator in the presence of 5% CO_2_ for 24h – 72h. Agar gels were diluted with 1 mL PBS, melted at 85°C for 3 minutes, and centrifuged at 40°C (5 minutes at 1800g) to pellet the cells. Cells were washed once with PBS and processed for cell cycle analysis as described ^26^. Agarose with control cells was melted and processed, as indicated, immediately after the cooling step.

Cell viability and ROS and glutathione intracellular content of cells cultured in liquid medium were assessed by flow cytometry analysis of cells stained for 30 minutes in the dark with 0.025 µM calcein-AM (ThermoFisher Scientific, C3099) at room temperature, 50 µM Dichlorodihydrofluorescein-diacetate (DCFDA, ThermoFisher Scientific, D399) at 37°C, and 10 µM monochlorobimane (ThermoFisher Scientific, M1381MP) at 37°C, respectively. For analysis of cells embedded in soft agar, gels were prepared as for cell cycle analysis and kept at 37°C in a humidified incubator in the presence of 5% CO_2_ for 24h – 72h. When appropriate, liquid medium and soft agar were supplemented with 50 µM BSO. Agar gels were mixed with 1 mL PBS by vigorous pipetting before the addition of calcein-AM or mBcl to the same concentration as indicated for cells cultured in liquid medium and incubated for 30 minutes in the dark at the temperature indicated for those cells. A FACS Canto II flow cytometer (BD Biosciences) was used for the analysis employing Kaluza software (Beckman Coulter, Brea, CA, USA).

Splenocytes from λ-MYC mice and wild-type littermates were isolated as previously described ^27^ and stained with APC-conjugated rat anti-mouse CD45R/B220 (ref 561880) and PE-conjugated rat anti-mouse CD43 (ref 561857) antibodies, both from BD Biosciences, before flow cytometry analysis with a FACS Canto II flow cytometer (BD Biosciences) employing Kaluza software (Beckman Coulter)

### Glutathione and ROS staining of primary T and B cells

Glutathione and ROS staining of human T and B cells was performed on cryopreserved or freshly isolated mononuclear cells from peripheral blood or bone marrow samples. Cells were first stained with 50 µM DCFDA or 10 µM mBcl, as described above, followed by a washing step with PBS and a surface staining of 30 minutes on ice with combinations of the following monoclonal antibodies: PE-conjugated anti-CD10 (HI10α), PerCP-conjugated anti-CD3 (SK7), APC-conjugated anti-CD19 (SJ25C1), PE-conjugated anti-CD5 (L17F12), PE-conjugated anti-CD25 (PC61), all from BD Biosciences. After another washing step, samples were immediately acquired on a BD FACSCanto™ II with FACSDIVA software (BD).

### Gene expression analysis

Total RNA was extracted using RNeasy (Qiagen; Venlo, Netherlands) following manufacturer’s instructions. cDNA was prepared from total RNA and used for gene expression by real-time quantitative RT–PCR (qPCR) as described ^27^ using TaqMan Gene Expression Assays (ThermoFischer Scientific) specific for human *GCLC* (Hs00155249_m1) and *ACTB* (Hs01060665_g1). *ACTB* was chosen as a control gene on the basis of its homogeneous expression in non-transduced and control-transduced cells. Each gene expression experiment was performed at least 3 times and calculations were made from measurements of 3 replicates of each sample.

### Statistical analysis

The numbers of animals used are described in the corresponding figure legends. All experiments were carried out with at least three biological replicates. Experimental groups were balanced in terms of animal age, sex, and weight. Animals were caged together and treated in the same way. No randomization was used to allocate animals to experimental groups and investigators were not blinded to the group allocation during experiments or outcome assessments. GraphPad Prism software 9.4.1. was used for the analysis. Variance was comparable between groups throughout the manuscript. Differences were analyzed by one-way or two-way analysis of variance (ANOVA) with Bonferroni post-test, Student t-test, multiple t-test with Holm-Sidak method, Mann-Whitney test, or Mantel-Cox test, as appropriate. Differences were considered significant at p<0.05.

## Results

### Soft gels selectively support the survival of a subset of lymphoid tumor cells

Growth rates of LCLs (X50-7 and JY) and lymphoma B cells (DG-75, BL-2, Toledo) in liquid medium are nearly identical^17^. In line with previous findings^16,17,23^, lymphoma B cells form colonies in soft-agar hydrogels, unlike LCLs (Figure S1). PBLs require proper stimulation and IL-2 for in vitro proliferation^28^. To understand why non-tumor lymphoid cells do not grow in soft hydrogels, we cultured activated PBLs, LCLs, and lymphoma cells in such gels for 24-72 hours and then analyzed their cell cycle and viability using flow cytometry. We established the experimental conditions to recover cells embedded in the hydrogel and stain them with propidium iodide. Within just 24 hours in the soft gel, 40-50% of PBLs, JY, and X50-7 non-tumor cells underwent apoptosis, as indicated by an increase in the sub-G_0_/G_1_ population, and this effect was exacerbated after 48 hours (Figures 1A,1B). In contrast, <20% of Burkitt’s lymphoma cells (DG-75 and BL-2) or Diffuse Large B-Cell lymphoma cells (Toledo) entered apoptosis after 24 hours (Figures 1A,1B). While DG-75 cells modestly accumulated in the G_2_/M phase of the cell cycle, JY exhibited a slight decrease in the S phase, and the cell-cycle distribution of PBLs and X50-7 remained largely unaffected (Figure 1C).

**Figure 1.**
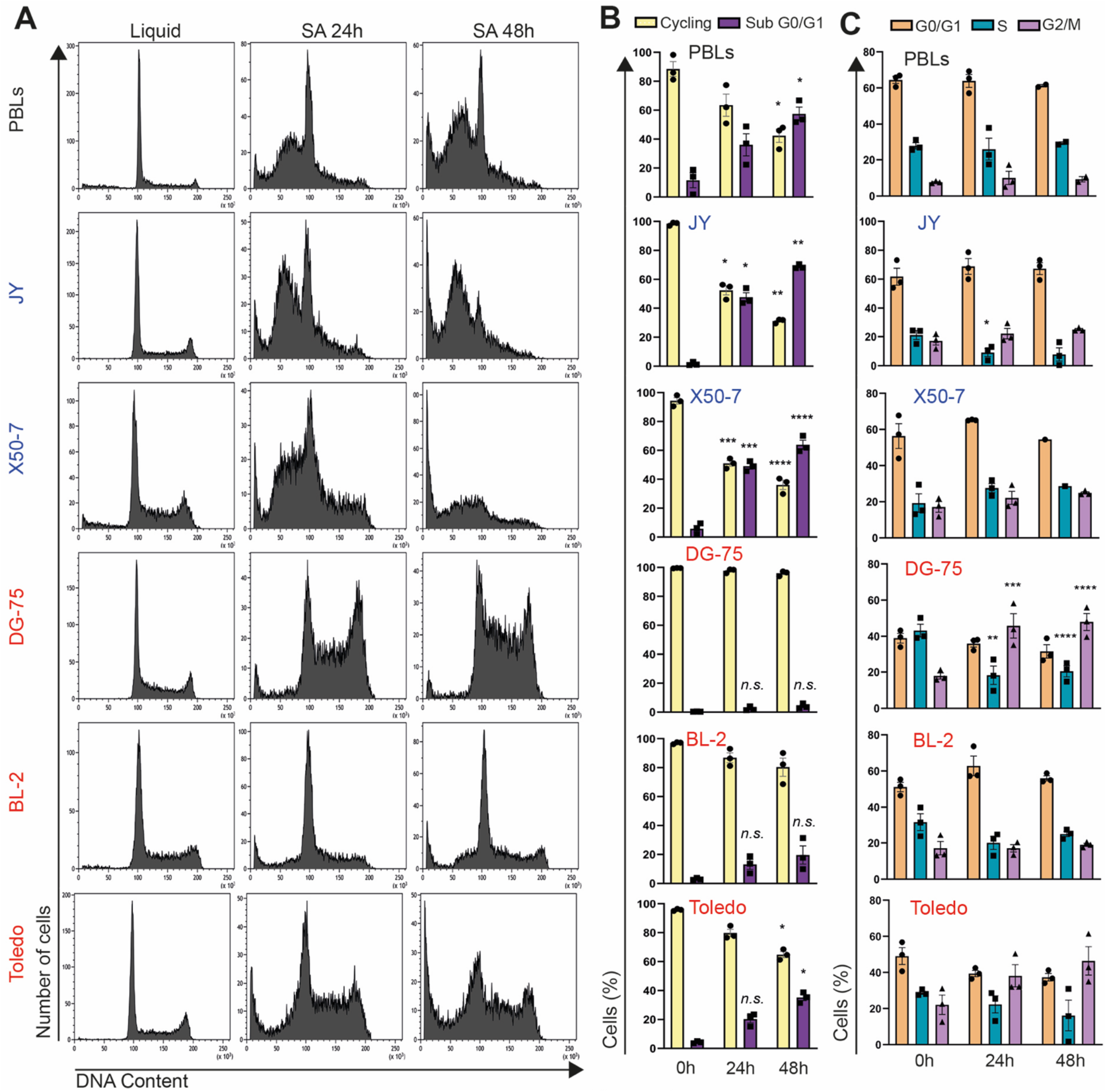
Cell death limits growth of lymphoid cells in soft gels. (**A**) Representative cell cycle profiles, (**B**) quantification of cycling cells (cells in G0/G1, S and G2/M) and apoptotic cells (SubG0/G1), and (**C**) quantification of cells in either G0/G1, S, or G2/M phase of the cell cycle in PBLs, LCLs (JY and X50-7), and lymphoma B-cells (DG-75, BL-2, and Toledo) grown in liquid culture or in soft-agar hydrogels (SA) for 24h or 48h. (B,C) Each data point denotes the value from an independent experiment and data in histograms are presented as mean ± s.e.m. *p<0.05, **p<0.01, ***p<0.001, ****p<0.0001 vs 0h; *n.s.*, non-significant; two-way ANOVA with Bonferroni post hoc test.

Flow cytometry analysis of cell size (FSC) and granularity (SSC) detected changes in cell viability (Figure S2), revealing a substantial decrease in viability of JY, X50-7, BL-2, and Toledo cells after 24 hours of culture in soft-agar hydrogels, with a milder reduction in DG-75 cells viability (Figure S3). The decline in viability among B-cell lymphomas was accentuated after 48 hours, and this effect was also observed in the Jurkat acute T-cell leukemia cell line (Figure S3). Similar results were obtained when analyzing cells stained with calcein-AM, a viability marker (Figure S4). Together, these findings strongly suggest that the limited growth of most lymphoid cells in soft hydrogels primarily results from cell death, rather than cell-cycle arrest, while a small subset of lymphoid tumor cells, likely TIC or TIC-like cells, maintain viability within the soft gel.

### Oxidative stress induces cell death in soft gels

Given that oxidative stress, an imbalance in ROS production and elimination leads to DNA damage and cell death^29^, we investigated whether cell culture in soft hydrogels increased ROS generation. Using flow cytometry, we analyzed cells stained with 2’,7’–dichlorofluorescein diacetate (DCFDA), a fluorescent dye for ROS detection. In liquid medium, LCLs and lymphoma cells showed similar intracellular ROS levels, with higher ROS in Jurkat cells (Figure 2A).

**Figure 2.**
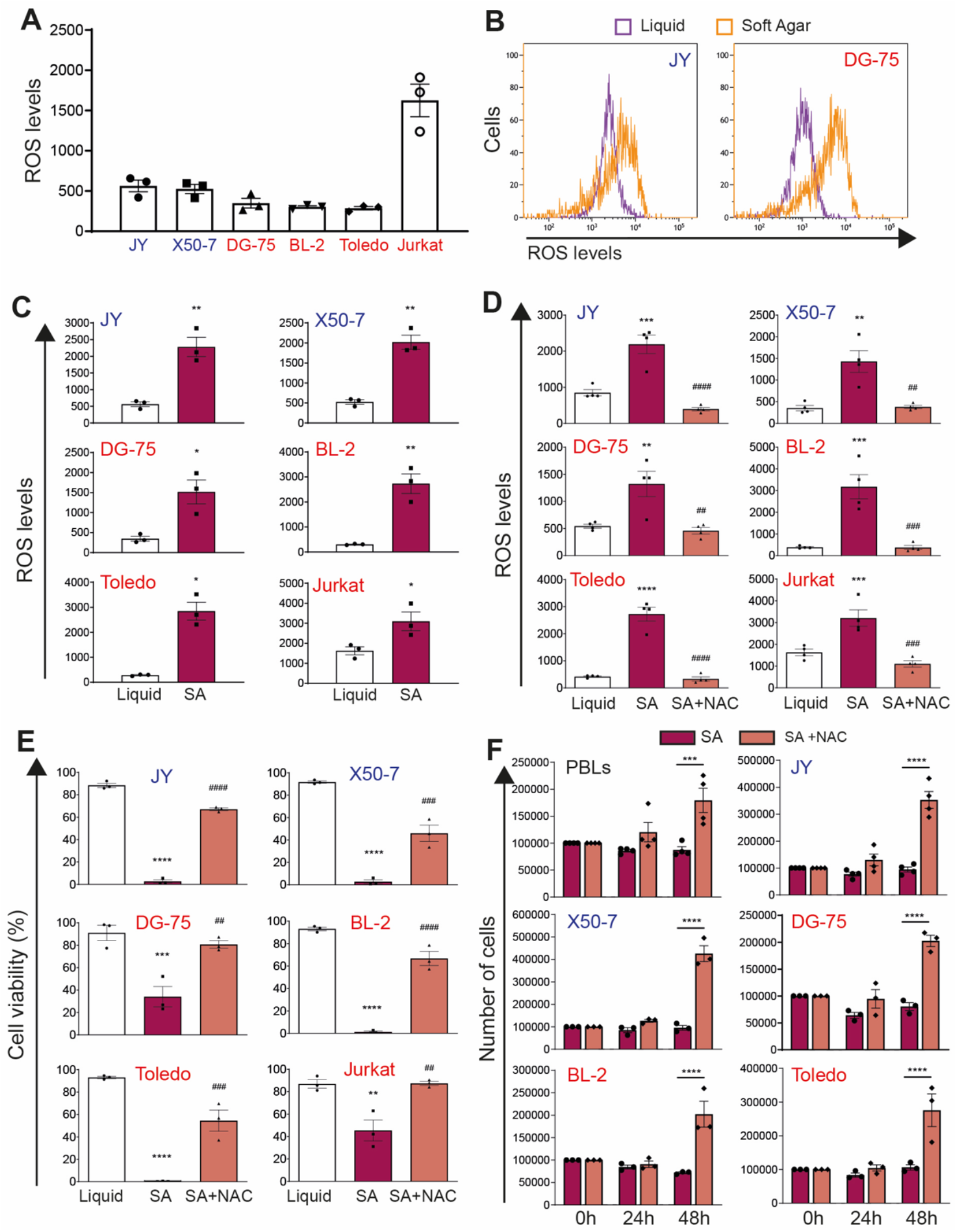
Increased oxidative stress mediates the lethality induced by cell culture in soft gels. Flow cytometry analysis of ROS staining with DCFDA in the indicated cells cultured in (**A**) liquid medium; (**B,C**) liquid medium or soft-agar hydrogels (SA) for 24h; and (**D**) liquid medium and SA for 72h in the presence of 5 mM N-acetylcysteine (NAC), as indicated. (**E)** Flow cytometry analysis of calcein staining in the indicated cells cultured either in liquid medium or in SA for 72h. (A,C,D,E) Each data point denotes the value from an independent experiment and data in histograms are presented as mean ± s.e.m. (C) *p<0.05, **p<0.01; Student t-test; (D,E) **p<0.01, ***p<0.001, ****p<0.0001 vs liquid; ^##^p<0.01, ^###^p<0.001, ^####^p<0.0001 vs SA 72h; one-way ANOVA with Bonferroni post hoc test. (**F**) Primary normal peripheral blood lymphocytes (PBLs) and JY, X50-7, DG-75, BL-2, and Toledo cells (10^5^ cells) were seeded in SA in the absence (SA) or presence of 5 mM NAC (SA + NAC) and recovered immediately after jellification (0) or after 24h and 48h of culture in this medium. Each data point denotes the number of cells recovered in an independent experiment and data in histograms are presented as mean ± s.e.m. ***p<0.001, ****p<0.0001; two-way ANOVA with Bonferroni post hoc test.

However, after 24 hours in soft-agar gels, all cell types exhibited a sharp rise in ROS levels (Figures 2B,2C). Addition of N-acetylcysteine (NAC), a potent antioxidant, substantially decreased ROS levels in cells cultured for 72 hours in these gels (Figure 2D), improved the viability of lymphoid tumor cells, LCLs, and PBLs from healthy donors in the hydrogel (Figure 2E) and even promoted rapid growth in soft-agar gels for these cell types (Figure 2F).

### Glutathione is required for lymphoid tumor cell growth in soft gels and xenografts

To neutralize ROS, cells induce various antioxidant systems, including the activation of the protein machinery that produces glutathione^30^, the most abundant antioxidant scavenger in all cells^31,32^. To investigate whether the survival of TIC-like cells in soft gels is mediated by ROS neutralization with glutathione, we assessed intracellular glutathione levels in lymphoid tumor cells and LCLs. Flow cytometry analysis of cells stained with mBcl, a fluorescent dye for glutathione detection, revealed similar intracellular glutathione levels in LCLs and lymphoma/leukemia cells in liquid medium (Figure 3A). However, when cultured in hydrogels for 24 hours, lymphoid tumor cells increased glutathione levels, while levels remained unchanged or decreased in LCLs (Figures 3B,3C).

**Figure 3.**
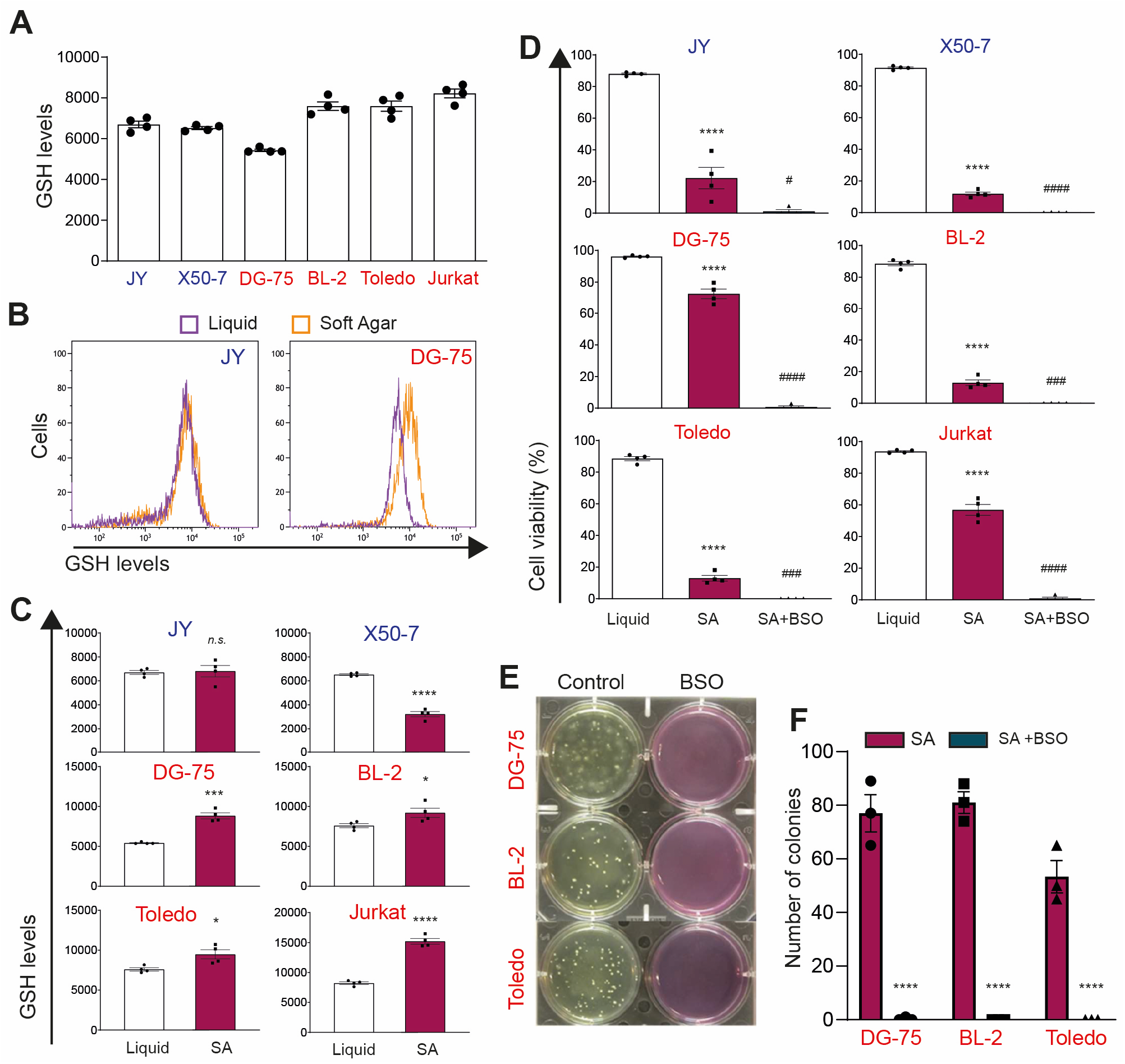
Glutathione overproduction mediates survival and growth of TIC in soft gels. Flow cytometry analysis of glutathione (GSH) staining with mBcl in the indicated cells cultured in (**A**) liquid medium and (**B,C**) liquid medium or soft-agar hydrogels (SA) for 24h. (**D**) Rate of living cells cultured 24h in liquid medium, SA, or SA supplemented with the GCL inhibitor BSO (50 µM), as determined by flow cytometry analysis of cell size and granularity. (**E**) Images of representative wells containing the indicated cells grown in SA or SA supplemented with 50 µM BSO for 21 days and (**F**) number of colonies per well. (A,C,D,F) Each data point denotes the value from an independent experiment and data in histograms are presented as mean ± s.e.m. (C) *p<0.05, ***p<0.001, ****p<0.0001, n.s., non-significant vs Liquid, Student t-test. (D) ****p<0.0001 vs Liquid; ^#^p<0.05, ^###^p<0.001, ^####^p<0.0001 vs SA; one-way ANOVA with Bonferroni post hoc test. (F) ****p<0.0001 vs SA, two-way ANOVA with Bonferroni post hoc test.

The rate-limiting reaction in glutathione synthesis is catalyzed by glutamate cysteine ligase (GCL)^33^. To determine glutathione contribution to TIC-like survival in soft gels, we treated lymphoid tumor cells with DL-Buthionine-sulphoximine (BSO), a pharmacological inhibitor of GCL, and found that this treatment markedly decreased glutathione levels (Figure S5A) and induced ROS accumulation (Figure S5B) in LCLs and lymphoid tumor cells in liquid culture without substantially decreasing their viability (Figure S5C). In contrast, BSO treatment blunted the viability of lymphoid tumor cells in soft-agar hydrogels (Figures 3D) and blocked colony formation (Figures 3E,3F).

GCL comprises a catalytic (GCLC) and a modifier (GCLM) subunit^33^. To further confirm the contribution of glutathione to TIC-like cells survival in soft gels, we silenced *GCLC* in DG-75 lymphoma cells with shRNA-encoding lentivirus. Among the candidate shRNAs targeting *GCLC,* sh-62, sh-65, and sh-86 demonstrated high knockdown efficacy compared to a non-targeting shRNA (sh-Ctl) in these cells (Figure 4A). While *GCLC* silencing markedly decreased glutathione levels in DG-75 cells without affecting their viability in liquid medium, it sharply decreased glutathione accumulation and cell viability, and nearly blocked their growth in soft-agar hydrogels (Figures 4B-4F). To evaluate the role of glutathione in the *in vivo* tumorigenicity of lymphoma cells, we subcutaneously inoculated DG-75 cells transduced with sh-Ctl or *GCLC*-specific sh-65 and sh-86 in immunodeficient NOD-SCID mice. As expected, control-transduced DG-75 cells readily elicited growth of large tumors in these mice (Figure 4G). Conversely, *GCLC* silencing markedly impaired tumor growth (Figure 4G). Similar results were observed in Toledo Diffuse Large B-Cell lymphoma cells. *GCLC* silencing decreased glutathione accumulation without affecting viability in liquid medium (Figures S6A-S6C), but markedly diminished glutathione levels (Figure S6D), sharply decreased viability (Figure S6E), and blocked cell growth in soft-agar hydrogels (Figure S6F). Together, these results emphasize the critical role of glutathione in the growth of TIC-like cells in soft gels and tumorigenicity.

**Figure 4.**
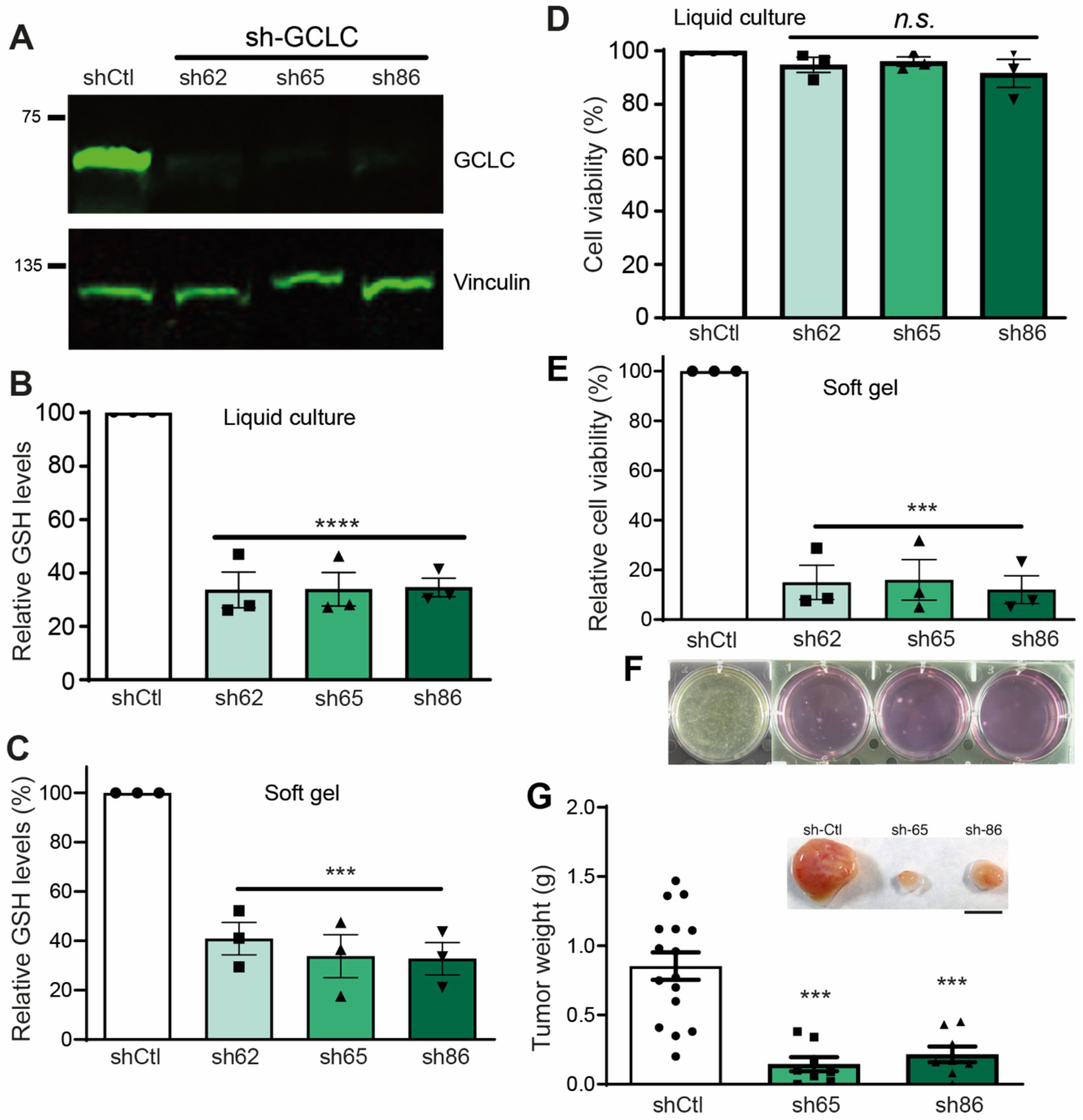
*GCLC* silencing impairs lymphoma cells survival and growth in soft gels and lymphomagenesis *in vivo*. DG-75 cells were transduced with lentivirus encoding a puromycin-inactivating protein and either a control shRNA (shCtl) or the *GCLC*-specific shRNAs sh-62, sh-65, or sh-86. Transduced cells were selected by culture for >5 days in liquid medium containing 2 μg/ml puromycin and then further cultured in soft-agar hydrogels (SA). (**A**) Immunoblot analysis for GCLC and Vinculin (loading control) expression in protein extracts from DG-75 cells transduced with lentivirus expressing the indicated shRNAs. Flow cytometry analysis of the staining of the indicated transduced cells with (**B, C**) mBcl or (**D, E**) calcein after 24h of culture in liquid medium (B, D) or soft agar (C, E). Each data point denotes the value from an independent experiment and data in histograms are presented as mean ± s.e.m. ***p<0.001 one-way ANOVA with Bonferroni post hoc test. (**F**) Images of representative wells containing the indicated transduced DG-75 cells grown in SA for 21 days. (**F**) Image of representative tumors and weight of tumors in NOD-SCID mice subcutaneously inoculated with DG-75 cells transduced with shCtl (n = 16 mice), sh-65 (n = 8 mice), or sh-86 (n = 8 mice). Each data point denotes the tumor of an individual mouse, and the horizontal bars denote the mean (long bar) and s.e.m.; ***p<0.001 vs sh-Ctl; one-way ANOVA with Bonferroni post hoc test. Bar, 1 cm.

### Glutathione and ROS dysregulation in tumor cells from lymphoma/leukemia patients

We aimed to investigate the role of the ROS-glutathione pathway in human cancer by quantifying intracellular ROS and glutathione levels in tumor B cells from B-cell lymphoma/leukemia patients and non-tumor B cells from healthy donors, with non-tumor T lymphocytes as controls. We found that the amount of ROS and glutathione in B lymphocytes relative to T lymphocytes from the same subject was markedly higher in B-cell lymphoma or leukemia patients than in healthy donors (Figure 5), suggesting that human lymphoma/leukemia cells may increase glutathione production to neutralize the toxic effects of increased ROS production.

**Figure 5.**
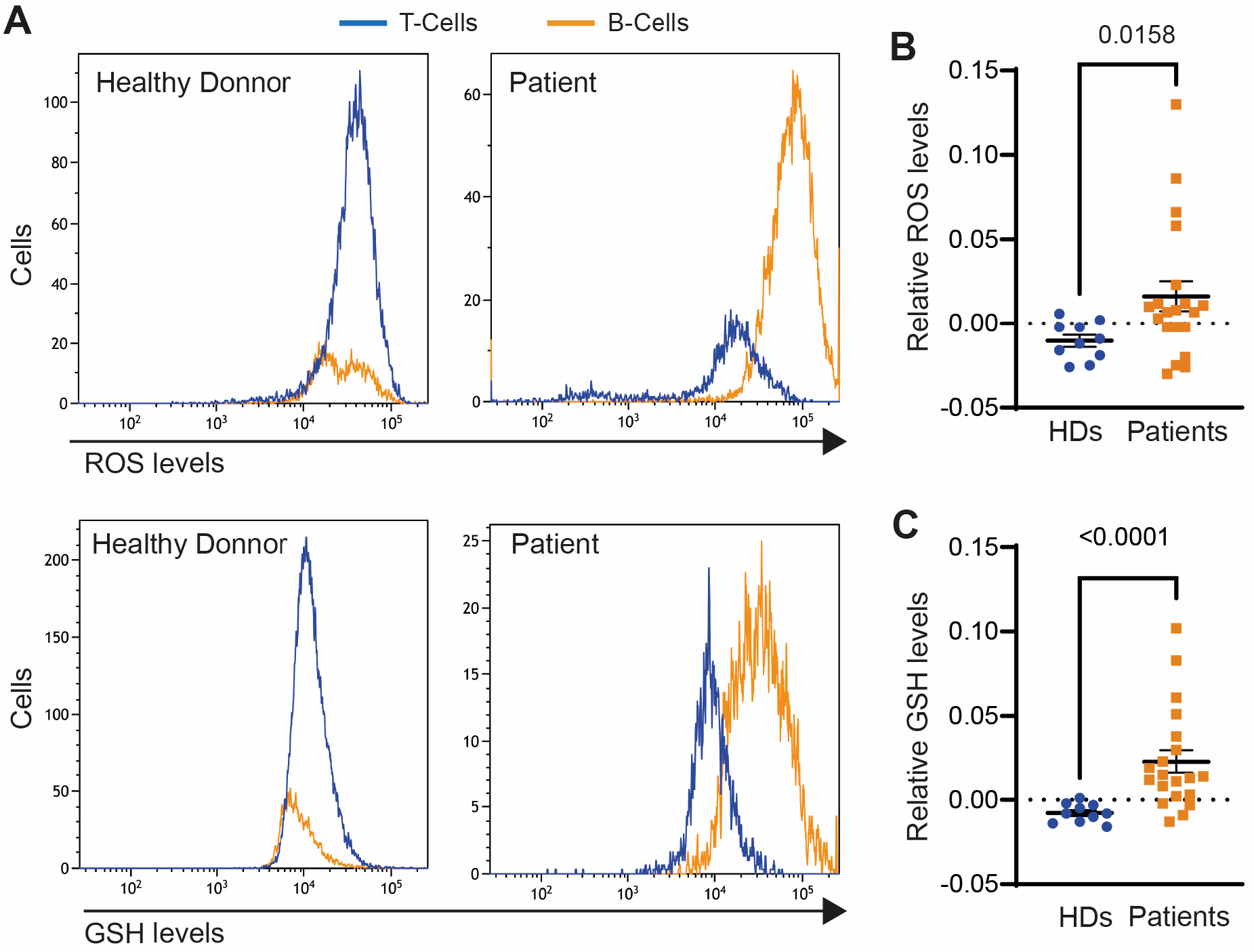
Lymphoid tumor cells contain higher glutathione and ROS levels than non-tumor lymphoid cells. Total blood cells from 10 healthy donors (HD) and 6 acute lymphoblastic B-cell leukemia, 5 chronic lymphoblastic B-cell leukemia, 2 follicular lymphoma, 2 mantle cell lymphoma, 2 splenic marginal zone lymphoma, 1 small lymphocytic lymphoma/chronic lymphocytic leukemia, 1 diffuse large B-cell lymphoma, and 1 hairy-cell leukemia patients were labeled with anti-CD3 and anti-CD19, to identify T and B lymphocytes respectively, and with DCFDA or mBcl as indicators of intracellular ROS and glutathione (GSH), respectively. (**A**) DCFDA and mBcl staining of a representative HD and patient. The ratio of (**B**) ROS and (**C**) glutathione (GSH) levels in B cells relative to T cells of the same subject is shown. Each data point denotes an individual, and the horizontal bars denote the mean (long bar) and s.e.m. Differences were analyzed by Mann-Whitney test (p values are shown).

### Glutathione overproduction is required for tumor growth in females of a lymphoma mouse model

To assess the translational potential of our findings, we investigated the impact of glutathione-mediated ROS neutralization on lymphomagenesis in a preclinical B-cell lymphoma mouse model, C57BL/6N-Tg(Igl-MYC)3Hm (1-Myc mice)^25^. Lymphoma development in these mice becomes visually evident and palpable around 3 months of age, with >80% of both male and female mice displaying lymphoma signs before 6 months of age (Figure 6A). Necropsy confirmed splenomegaly and lymph node enlargement (Figures 6B,6C). Flow cytometry analysis of blood cells and splenocytes from λ-Myc mice demonstrated increased intracellular glutathione levels compared to age-matched control littermates (Figure 6D), indicating a potential requirement for ROS neutralization through glutathione overproduction in lymphomagenesis in vivo.

**Figure 6.**
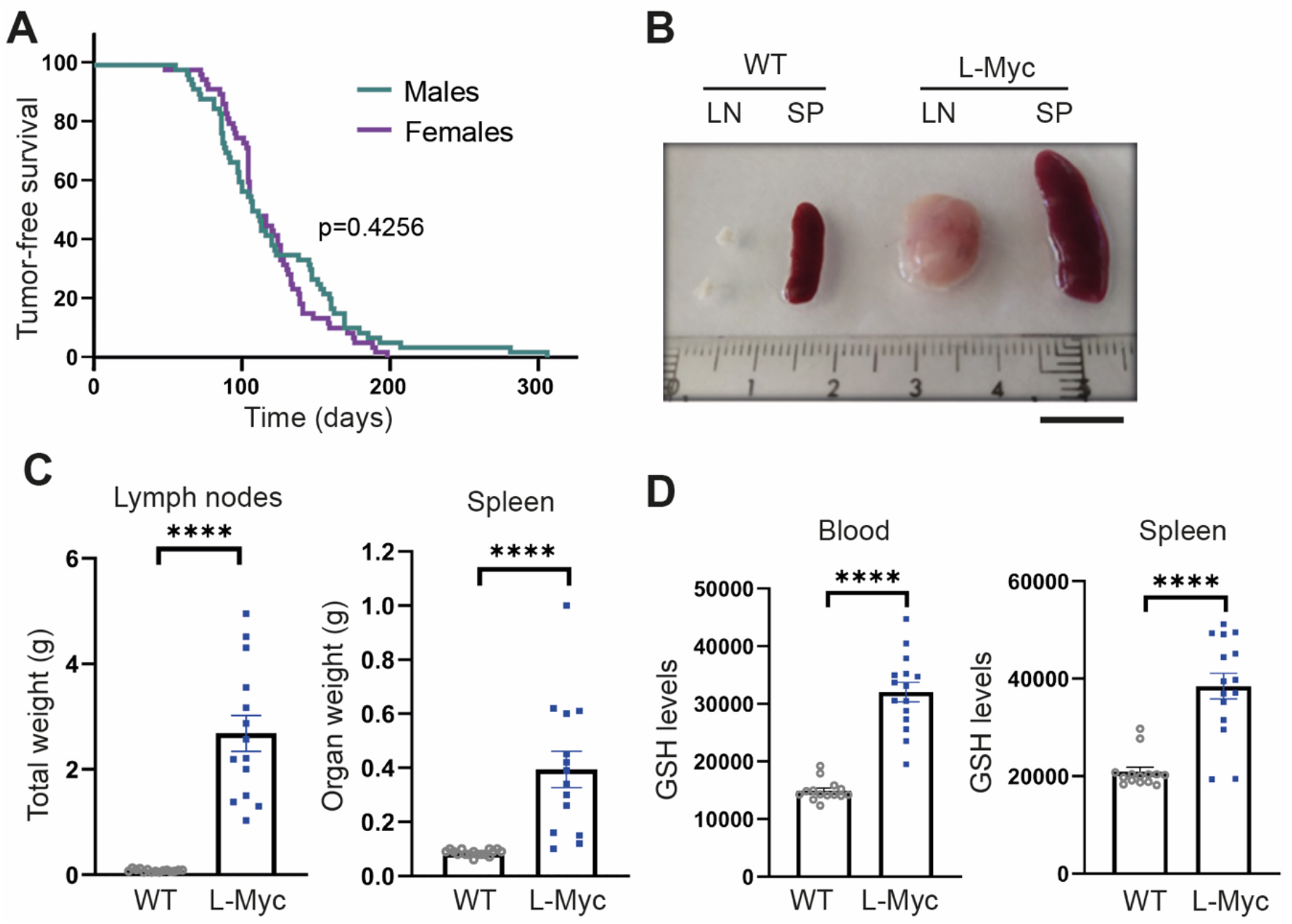
Increased glutathione levels in tumor tissues of a mouse model of B-cell lymphoma. (**A**) Tumor-free survival curve of 60 male and 60 female λ-Myc mice. Differences were analyzed by Mantel-Cox test; the p value is indicated. (**B**) Representative images and (**C**) spleen weight and aggregated weight of lymph nodes from each of 14 λ-myc mice and 17 WT littermates. Bar, 1 cm. (**D**) Flow cytometry analysis of mBcl staining (GSH levels) of blood cells and splenocytes from 15 λ- Myc and 14 WT littermate mice. Each data point denotes an individual and data in histograms are presented as mean ± s.e.m. ****p<0.0001; Student t-test.

To examine this hypothesis and evaluate the therapeutic prospects of pharmacological GCL inhibition, we treated λ-Myc mice with BSO. As previously reported^34^, an increase in B220^+^/CD43^+^ double-positive tumor cells was observed in the spleen of these mice at 3 weeks of age (Figure 7A). We therefore initiated treatment with BSO in the drinking water for both male and female 1-Myc mice at P21 and maintained it until 8 months of age, unless humane endpoints were reached. While male mice exhibited minimal changes in tumor formation, treatment of female mice sharply delayed tumor growth (Figure 7B). Notably, this treatment markedly decreased glutathione levels in splenocytes of male and female mice (Figure 7C). Importantly, long-term BSO treatment did not substantially affect mouse body weight (Figure 7D), suggesting the lack of noticeable toxic effects.

**Figure 7.**
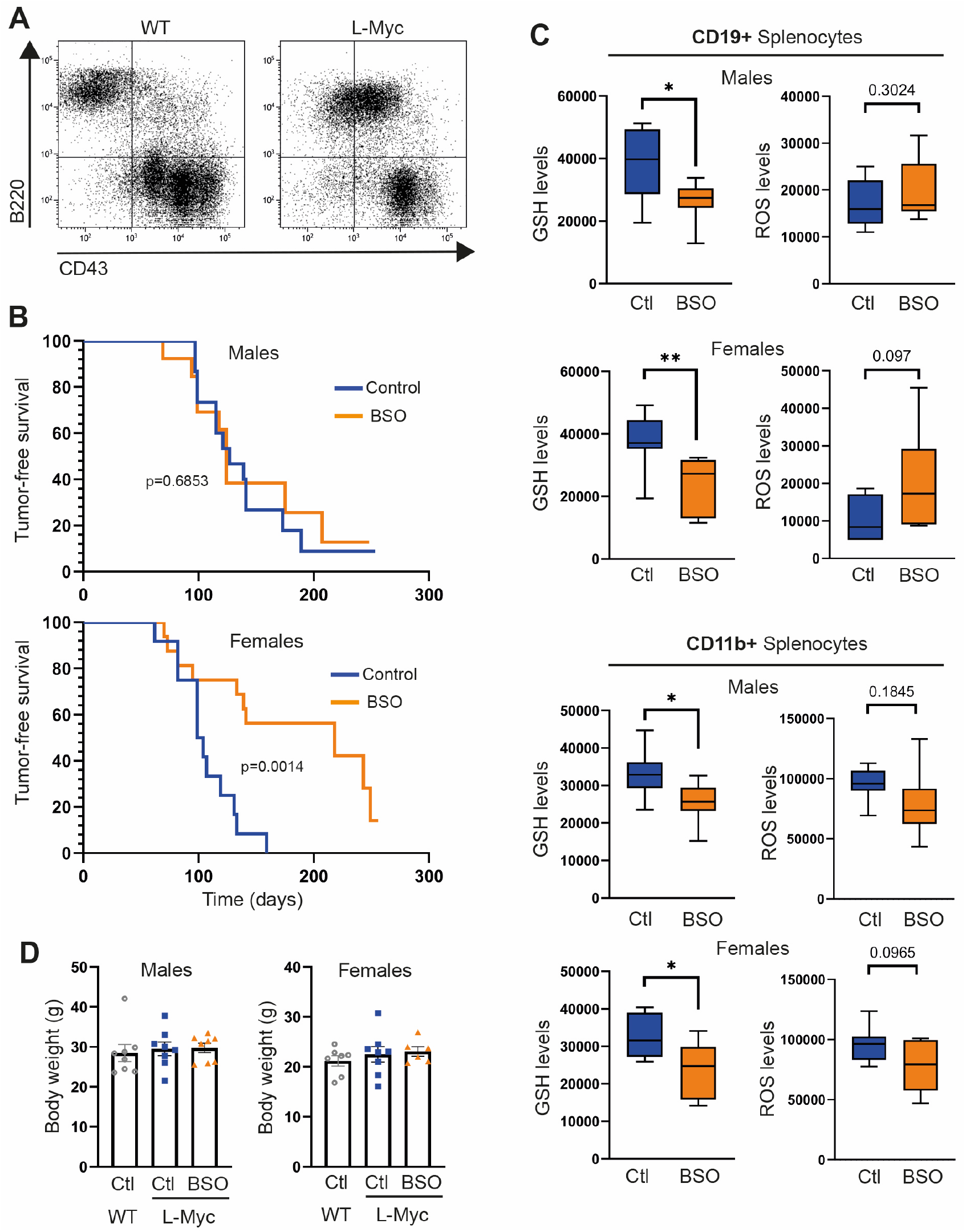
Dimorphic effect of glutathione synthesis inhibition on lymphoma growth in a mouse model of B-cell lymphoma. (**A**) Representative B220/CD43 double staining of splenocytes from 21-day-old λ-Myc mice. (**B**) Tumor-free survival curve of 15 male and 12 female untreated (Control) λ-Myc mice and 13 male and 16 female λ-Myc mice treated with BSO in the drinking water (20 mM). Differences were analyzed by Mantel-Cox test (p values are shown). (**C**) End-of-experiment flow cytometry analysis of mBcl (GSH levels) and DCFDA (ROS levels) staining of CD19+ and CD11b+ splenocytes from 8 male and 7 female untreated (Ctl) λ-Myc mice and 8 male and 7 female λ-Myc mice treated with BSO. Data are presented as box-and-whisker plots, with 75th and 25th percentiles; bars represent maximal and minimal values. *p<0.05, **p<0.01, n.s., non-significant by multiple t-test with Holm-Sidak method. (**D**) End-of-experiment body weight of 8 male and 7 female untreated (Ctl) λ-Myc mice and 8 male and 7 female λ-Myc mice treated with BSO. Each data point denotes an individual and data in histograms are presented as mean ± s.e.m.

## Discussion

To discover therapeutic strategies that prevent tumor recurrence, we have focused on a unique trait of TICs, their capacity to grow in soft gels. Since this characteristic strongly correlates with tumorigenicity in animals^12^, we hypothesized that interventions inhibiting TIC growth in soft gels could also impede tumor development *in vivo*. Indeed, our study reveals that inhibiting glutathione synthesis effectively halts the growth of lymphoma cells in soft agar hydrogels and hampers lymphomagenesis in a mouse lymphoma model. We have also shown that primary lymphoma/leukemia cells from this mouse model and B-cell lymphoma/leukemia patients accumulate higher intracellular glutathione levels than non-tumor lymphocytes. This suggests that glutathione overproduction may be indispensable for the survival and proliferation of primary tumor cells, making the inhibition of glutathione production a promising therapeutic avenue for lymphoid tumors.

Conventional cancer treatment targets are often related to cell proliferation, likely due to their identification via comparisons between actively proliferating cancer cells and slowly dividing or non-dividing normal diploid non-tumor cells. To circumvent this potential bias, we chose to compare lymphoma/leukemia cells with LCLs, as both cell types grow indistinguishably in liquid culture^17^. While these lymphoid cell lines nearly double their numbers every 24 hours in liquid culture, a critical distinction emerges in soft-agar hydrogels: LCLs do not grow, while a minority of lymphoid tumor cells form colonies in these soft gels^17,23^.

We have previously identified other genes required for lymphoma cell growth in soft gels and lymphomagenesis. Indeed, we demonstrated that silencing *E2F1* or forcing *E2F4* expression inhibited growth in soft gels. However, these manipulations on E2F factors also inhibited cell cycle progression in liquid medium^16,17^, revealing their involvement in broader cellular processes beyond TIC-specific functions. We also discovered that *CDCA7* silencing specifically inhibits cell growth in soft gels without inhibiting proliferation in liquid culture^23^. *CDCA7* silencing additionally hampered tumor cell migration and invasion^35^, strongly suggesting that CDCA7 inhibition could be of therapeutic interest for lymphoid tumors. However, the connection between CDCA7 mutations and a rare disease characterized by immunodeficiency, centromeric instability, and facial anomalies^36^ precludes the systemic administration of CDCA7 inhibitors as a lymphoma treatment.

To the best of our knowledge, the reason non-tumor lymphoid cells do not grow in soft gels remained unknown. Similar to LCLs, non-tumor epithelial and mesenchymal cells do not grow in soft-agar hydrogels, whereas a minor fraction of their tumor counterparts grows in these gels^13,15^. This phenomenon has been attributed to the reliance of non-tumor epithelial and mesenchymal cells on a rigid surface for survival^13^. However, the viability of non-tumor lymphoid cells in the circulatory system indicates that anchorage to a rigid surface is not required for their survival. Therefore, the growth disparity in soft gels may stem from a distinct mechanism. Our findings suggest that some lymphoid cells accumulate in G_0_/G_1_ or G_2_/M, while others experience minimal alteration cell cycle disruption. In contrast, increased cell death was consistently observed across various lymphoid cell types, including tumor cells, underscoring that, in soft gels, cell death, rather than cell cycle arrest, impedes lymphoid cell growth.

The inhibition of cell death by the anti-oxidant NAC supports the notion that oxidative stress causes the death of lymphoid cells in soft gels. Given that GCL pharmacological inhibition or silencing effectively blocks the growth of any lymphoid tumor cell in soft gels, including TIC-like cells, we propose that the minority of tumor cells thriving in soft gels counteracts ROS-induced cell death by boosting glutathione production. Of note, NAC not only prevents death of non-tumor lymphoid cell death in soft gels but also enables their growth in these conditions. Therefore, caution is advised regarding the administration of high antioxidant doses in diets because an excess might potentially promote overgrowth in antigen-stimulated lymphoid cells. Moreover, as >90% of adults harbor quiescent EBV-transduced B cells^37^, high doses of antioxidants might potentially trigger uncontrolled growth in these cells. This hypothesis aligns with reports of EBV presence in various human cancers^38^.

Neutralizing oxidative stress with glutathione not only supports lymphoid tumor cells proliferation in soft gels but also *in vivo*. This is evidenced by the inhibition of their growth in immunodeficient mice following lentivirus-mediated *GCLC* silencing. More importantly, pharmacological GCL inhibition impaired lymphomagenesis in a mouse model of B-cell lymphoma, although, for unknown reasons, this effect was observed only in female mice. These results are unexpected because females generally exhibit a more proficient defense against ROS than males ^29^.

TICs maintain lower intracellular ROS levels than other cancer cells in the tumor, largely due to ROS scavengers accumulation^6,7^. Since most chemotherapeutics increase intracellular ROS levels^39^, TIC resistance may arise from their high antioxidant content, including glutathione. Combining conventional chemotherapy to eliminate the bulk of tumor cells with drugs that inhibit glutathione synthesis to eradicate TICs might potentially achieve complete, relapse-free tumor remission. Moreover, this combination approach might allow for lower doses of conventional chemotherapy, reducing undesirable toxic effects. In this regard, long-term treatment with BSO did not substantially affect the weight of treated mice, suggesting limited undesired effects of this treatment.

## AUTHOR CONTRIBUTIONS

MRC conceived and designed the study with contributions from TI; AH-A. and OK performed most of the experiments with contributions from PM-N and EH-M.; AM-J and CM-C analyzed human samples; PF, MLT, and TI provided experimental support and ideas for the project. AH-A, OK and MRC analyzed the data with contributions from all other authors. MRC wrote the manuscript with contributions from xxx and TI. All authors approved the manuscript.

## ACKNOWLEDGEMENTS

The authors would like to thank D. Trono for plasmids and H.C. Morse III and the National Institute of Allergy and Infectious Dieseases (NIH) for donating C57BL/6N-Tg(IGL-MYC)3Hm mice. The CBMSO is supported by Consejo Superior de Investigaciones Científicas and Universidad Autónoma de Madrid, and is a Severo Ochoa Center of Excellence (grant CEX2021-001154-S) funded by MICIN/AEI/10.13039/501100011033. This work was supported by Fundación de la Asociación Española contra el Cáncer (AECC) grant PROYE20060CAMP to M.R.C., and grant PID2020-115218RB-I00 funded by MCIN/AEI/10.13039/501100011033, and Instituto de Salud Carlos III (CIBERNED) to T.I. O.K. held an FPI fellowship from the Spanish Ministerio de Educacion y Ciencia (BES-2014-069236).

## DATA AVAILABILITY

The datasets generated during and/or analyzed during the current study are available from the corresponding author on reasonable request.

