## Supplemental Figures for "Glutathione overproduction mediates lymphoma initiating cells survival and has a sex-dependent effect on lymphomagenesis"

### Supplementary Figures & legends

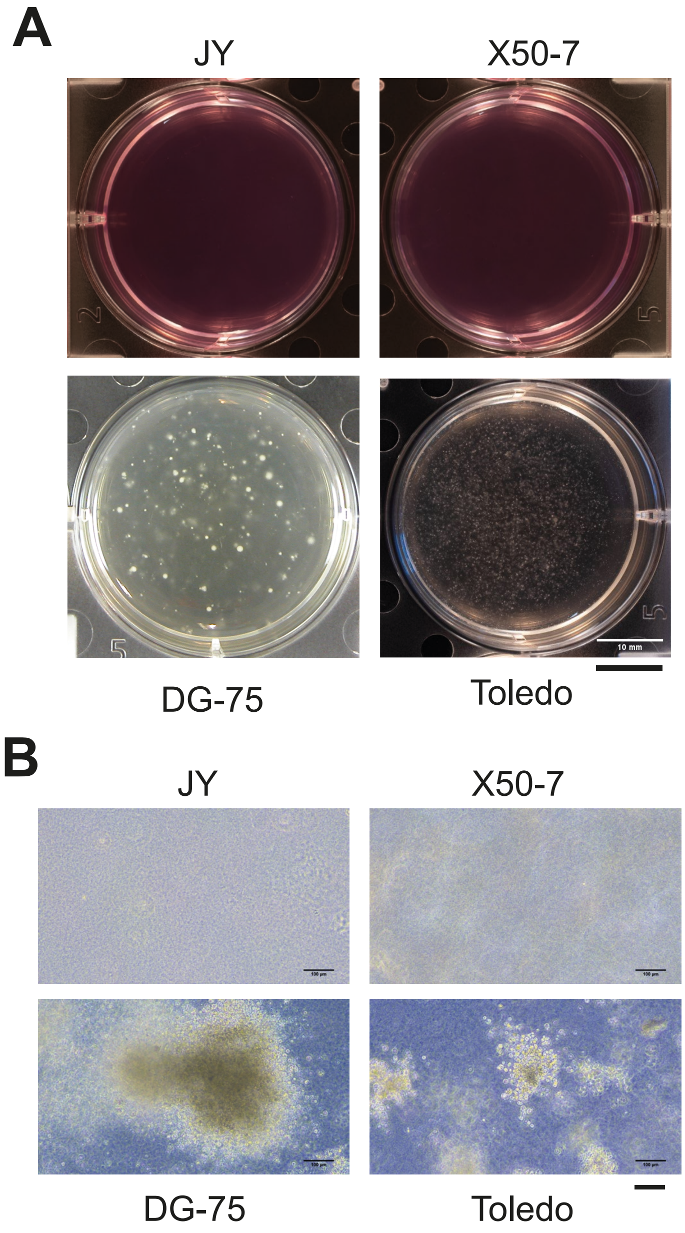


**Supplementary Figure 1. Lymphoma cells, but not LCLs, form colonies in soft gels.** (**A**) Macroscopic and (**B**) microscopic images of representative wells containing LCLs (JY and X50-7) or lymphoma cells (DG-75 and Toledo) grown in soft-agar hydrogels for 21 days. Bars, 10 mm (A) and 100 μm (B).

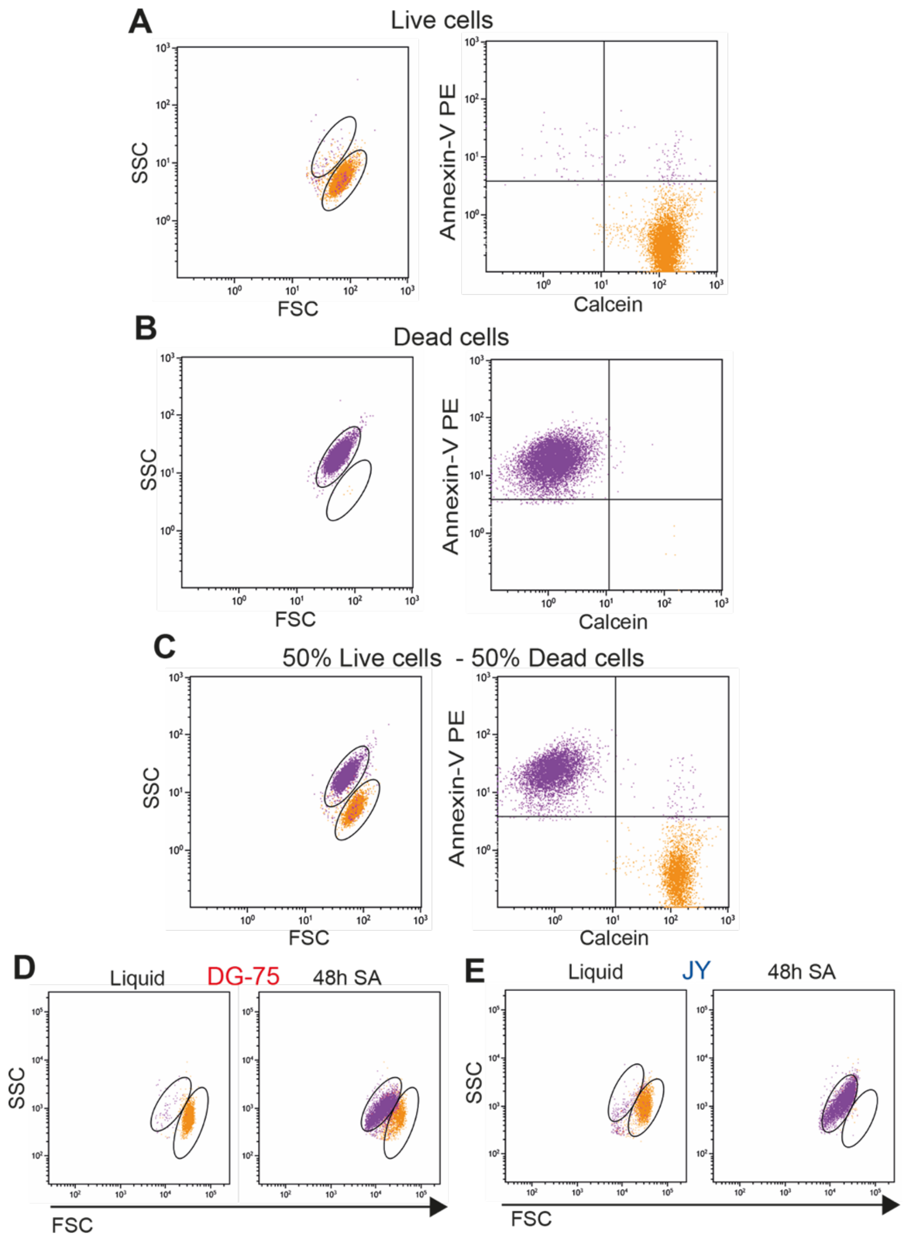


**Supplementary Figure 2. Cell size and granularity analysis by flow cytometry distinguishes live and dead cells.** Flow cytometry analysis of DG-75 cells incubated at 65ºC for 15 minutes or left untreated and stained with Calcein-AM and phycoerythrin-conjugated annexin V. Forward scatter (FSC) and side scatter (SSC) dot plots (left panels) and Annexin-V and Calcein dot plots (right panels) of (**A**) control DG-75 cells, (**B**) DG-75 cells incubated at 65ºC for 15 minutes, and (**C**) a 50% mixture of control and heated DG-75 cells. FSC/SSC dot plots of (**D**) DG-75 and (**E**) JY cells cultured in liquid medium or in SA for 48h.

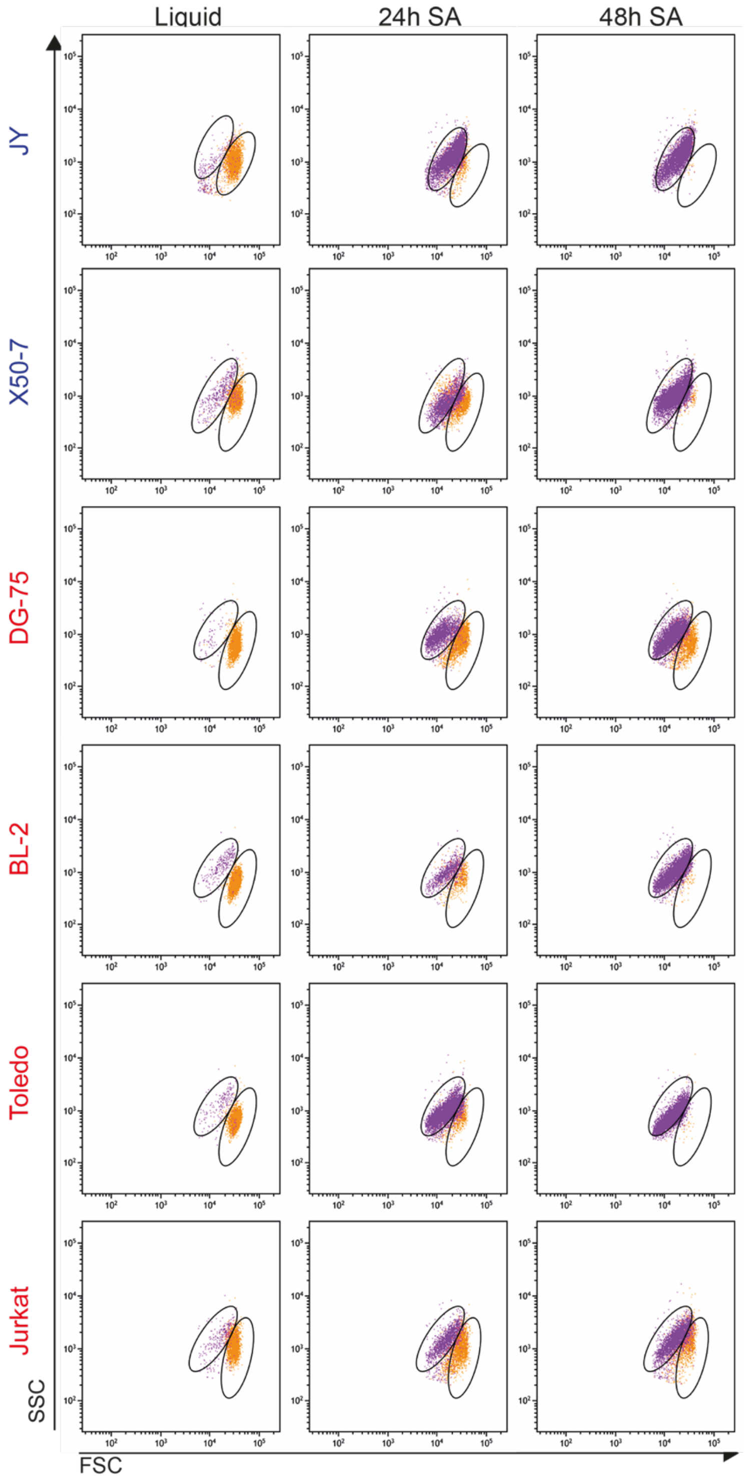


**Supplementary Figure 3. Cell culture in soft-agar gels induces cell death.** Flow cytometry analysis of cell size (FSC) and granularity (SSC) of the indicated cell lines cultured in liquid medium or in soft-agar (SA) for 24h or 48h. Live and dead cells populations are indicated.

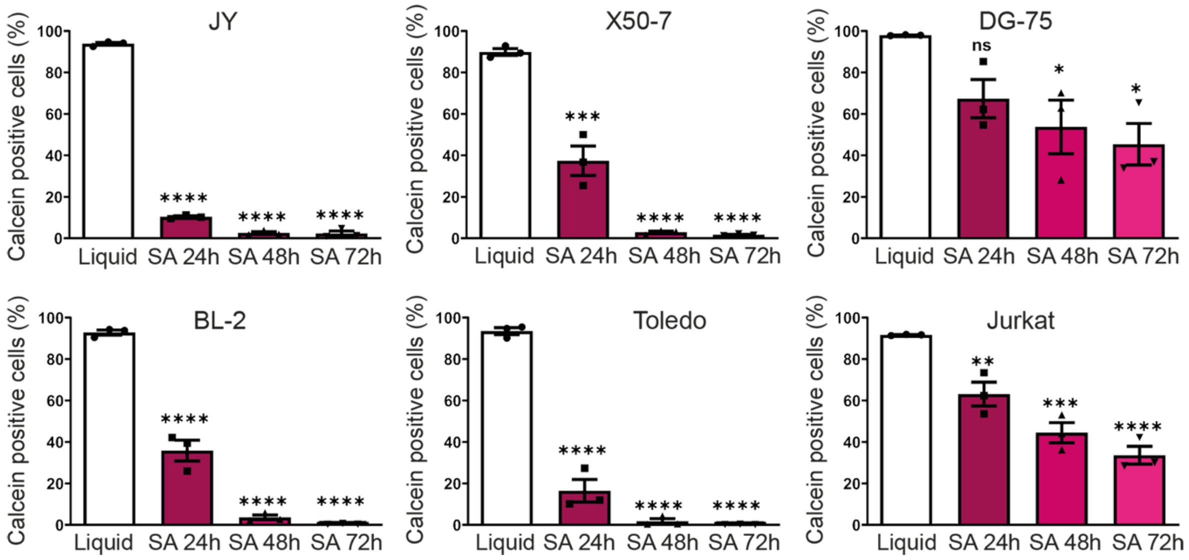


**Supplementary Figure 4. Cell culture in soft gels decreases cell viability.** Flow cytometry analysis of calcein staining of JY, X50-7, DG-75, Toledo, BL-2, and Jurkat cells cultured either in liquid medium or in soft-agar hydrogels (SA) for 24h, 48h, or 72h. Each data point denotes the value from an independent experiment and data in histograms are presented as mean + s.e.m. *p<0.05, **p<0.01, ***p<0.001, ****p<0.0001, n.s., non-significant; one-way ANOVA with Bonferroni post hoc test.

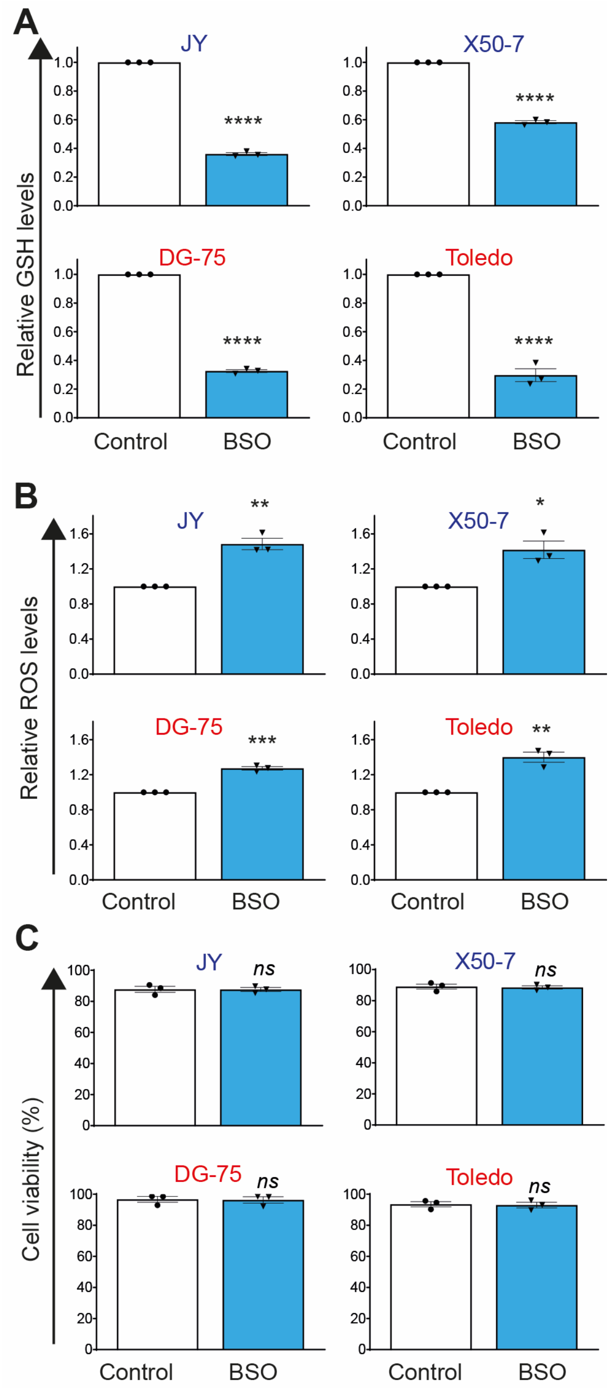


**Supplementary Figure 5. Glutathione synthesis inhibition increases oxidative stress but does not induce lethality in lymphoid cells grown in liquid medium**. The indicated cells were cultured for 24h in liquid medium in the absence (Control) or the presence of 50 µM BSO. Flow cytometry analysis of the staining of these cells with (**A**) mBcl or (**B**) DCFDA. (**C**) Rate of living cells as determined by flow cytometry analysis of cell size and granularity. Each data point denotes the value from an independent experiment and data in histograms are presented as mean ± s.e.m. *p<0.05, **p<0.01, ***p<0.001, ****p<0.0001, n.s., non-significant, Student t-test.

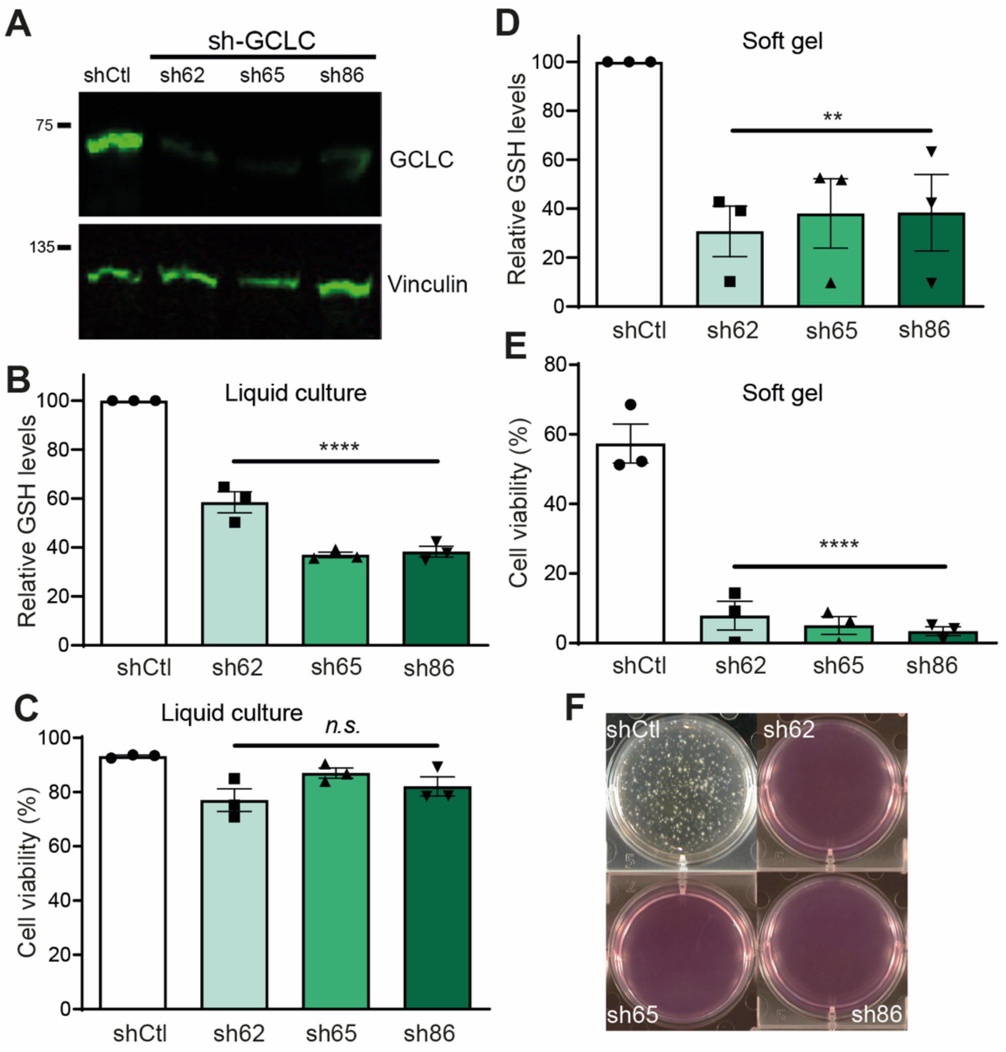


**Supplementary Figure 6. *GCLC* silencing impairs Toledo lymphoma cells survival and growth in soft gels**. Toledo cells were transduced with lentivirus encoding a puromycin-inactivating protein and either a control shRNA (shCtl) or the *GCLC*-specific shRNAs sh-62, sh-65, or sh-86. Transduced cells were selected by culture for >5 days in liquid medium containing 2 µg/ml puromycin and then further cultured (**A-C**) in liquid medium or (**D-F**) in soft-agar hydrogels. (A) Representative GCLC and Vinculin (loading control) immunoblot analysis of Toledo cells transduced as indicated (n = 3 independent cell batches per group). Flow cytometry analysis of (B,D) mBcl staining and (C,E) cell size and granularity of cells transduced as indicated after 24h of culture in liquid medium (B,C) or in SA (D,E). Each data point denotes the value from an independent experiment and data in histograms are presented as mean ± s.e.m. **p<0.01, ****p<0.0001, *n.s.*, non-significant, one-way ANOVA with Bonferroni post hoc test. (F) Images of representative wells containing the indicated transduced Toledo cells grown in SA for 21 days.
